# E2F1 induces a G0-G1 reentry transcriptional program without changing chromatin accessibility

**DOI:** 10.1101/2025.09.23.678145

**Authors:** Gerrald A. Lodewijk, Benjamin R. Topacio, Seungho Lee, Sayaka Kozuki, Znala Williams, Clara J. Han, Abolfazl Zargari, Tilini U. Wijeratne, Neda Bidoki, Silvart Arabian, Edward Wu, Eric Malekos, Joscha Weiss, Gerd A. Müller, Vanessa Jonsson, Nicolas H. Thomä, Seth M. Rubin, S. Ali Shariati

**Author notes:** Corresponding authors: Ali Shariati, Seth M. Rubin.

## Abstract

Quiescent cells actively repress cell-cycle genes via chromatin-based mechanisms to maintain a non-dividing state, yet remain poised to reenter upon stimulation. E2F1, a canonical activator of cell-cycle genes, is sufficient to induce reentry from quiescence, but how it overcomes chromatin-mediated repression remains unclear. Here, we show that inducible E2F1 expression triggers exit from quiescence and progression through the cycle without changes in chromatin accessibility, by harnessing regulatory elements with limited, pre-existing accessibility. Using time-resolved transcriptomics, we demonstrate that E2F1 induces an accelerated transcriptional program compared to serum. Unlike serum, which triggers broad chromatin remodeling, E2F1-induced activation occurs in a context of limited accessibility. ChIP-seq reveals that E2F1 directly binds target sites in quiescent cells to upregulate canonical genes. Biochemical reconstitution shows that E2F1 binds nucleosomes and accesses internal E2F sites within histone-wrapped DNA. These findings suggest that E2F1 can engage nucleosome-associated DNA and initiate transcription without major chromatin reorganization, redefining transcription factor–chromatin dynamics during cell fate transitions and establishing E2F1 as a potent regulator of cell-cycle reentry.

## Introduction

Many cells in the human body reside in a reversible nondividing quiescent state termed G0^1–4^. Cells enter quiescence when they do not receive the required signaling cues for cell-cycle progression such as growth factors. Quiescent cells can reenter the cell-cycle upon receiving growth factors, which results in activation of the Mitogen-Activated Protein Kinase Pathway (MAPK). Signaling through the MAPK pathway leads to increased activity of cyclin-dependent kinases (Cdks), at least in part through activation of Cyclin D expression. Cdks drive cell-cycle reentry by phosphorylation and inactivation of retinoblastoma (RB) family proteins, which in turn leads to activation of E2F transcription factors^5–10^. The E2F family of transcription factors contains eight homologs, which are further classified as activator and repressor E2Fs based on their effects on target gene expression^11,12^. E2F1 is known to be a transcriptional activator that binds to and induces expression of cell-cycle genes such as those encoding cyclins, cyclin-dependent kinases, and DNA replication machinery. Quiescent cells ensure fate stability against fluctuations in growth factor signaling by repression of these cell-cycle genes needed for S-phase entry. Repression depends on the RB protein and the multi-subunit transcriptional repressor complex, known as DREAM complex, which is composed of a repressor E2F, the RB paralog p130, and the MuvB core complex^7,8,13–15^. As a result, cell-cycle genes are repressed in quiescent cells to prevent activation via E2F transcription factors. However, it remains incompletely understood how different regulatory layers such as repressive chromatin marks that limit accessibility^4,16–19^, restricted binding of activator E2Fs at promoters^8,20^, and nucleosome positioning directed by MuvB^8,9,21^ contribute to this repression. It is thought that these specific chromatin lockdown mechanisms in quiescent cells ensure fate stability by restricting chromatin accessibility near cell-cycle genes, while allowing for reversibility to reenter the cell-cycle. How cells overcome restricted chromatin accessibility, establish multiple specific chromatin states with distinct capacities for gene activation, and utilize chromatin remodeling during exit from quiescence remain underexplored. Within this quiescent cell chromatin context, ectopic expression of E2F1 is sufficient to promote cell-cycle reentry independently of the upstream growth factor signals and MAPK activity^22,23^. Conversely, E2F1 deletion delays exit from quiescence suggesting that E2F1 is required for timely activation of a gene expression program required for the cell-cycle reentry^24^. Given that cell-cycle genes are actively repressed in quiescent cells, we were interested in studying how E2F1 on its own is able to activate expression of cell-cycle genes.

To better understand the chromatin-based mechanisms underlying quiescence exit, we systematically compared the dynamics, global gene regulatory mechanisms and chromatin state between E2F1-mediated cell-cycle reentry and growth factor-induced cell-cycle reentry. Our results show that E2F1 is able to rapidly promote quiescence exit by activating an accelerated gene expression program without global changes in chromatin accessibility. Unlike serum-induced cell-cycle reentry, we also find that E2F1-mediated gene transcription activation and subsequent cell division is not associated with increased chromatin accessibility. Notably, biochemical reconstitution demonstrates that E2F1 can directly bind to nucleosomes and can access internal E2F consensus sites within DNA wrapped by histones. Our findings support a model in which E2F1 promotes a gene expression program that allows cells to complete their cell division independent of the global changes in chromatin accessibility and likely through direct interaction with nucleosomes for a subset of genes.

## Results

### E2F1 expression alone is sufficient to exit quiescence and enter mitosis

To be able to control the expression of E2F1 in cells, we established a stable doxycycline (dox) inducible E2F1-Clover mouse fibroblast cell line (NIH-3T3) using a PiggyBac transposon system, and an empty-vector (EV) control (**Figure 1A**). We selected NIH-3T3 fibroblasts as our model system because they are well-established to enter a quiescent state in response to growth factor deprivation, making them suitable for studying cell-cycle reentry mechanisms^16,17^. To induce quiescence, cells were cultured in a starvation medium containing 0.25% serum for 48 hours. To initiate cell-cycle reentry, we added doxycycline (dox) to express E2F1 or switched the media to normal 10% serum levels (**Figure 1B**). Similar to previous reports^22,25^, we found that within 24 hours after dox-induction of E2F1, or serum addition, nearly 70% of cells entered the S-phase as indicated by incorporation of BrdU (**Figure 1C**). When we modulated E2F1 levels using increasing doses of doxycycline, we observed that quiescent cells reentered the cell-cycle only after reaching a specific threshold of E2F1 expression, suggesting that cells can detect defined E2F1 concentration thresholds for cell-cycle reentry (**Supplementary Figure 1A-B**).

**Figure 1.**
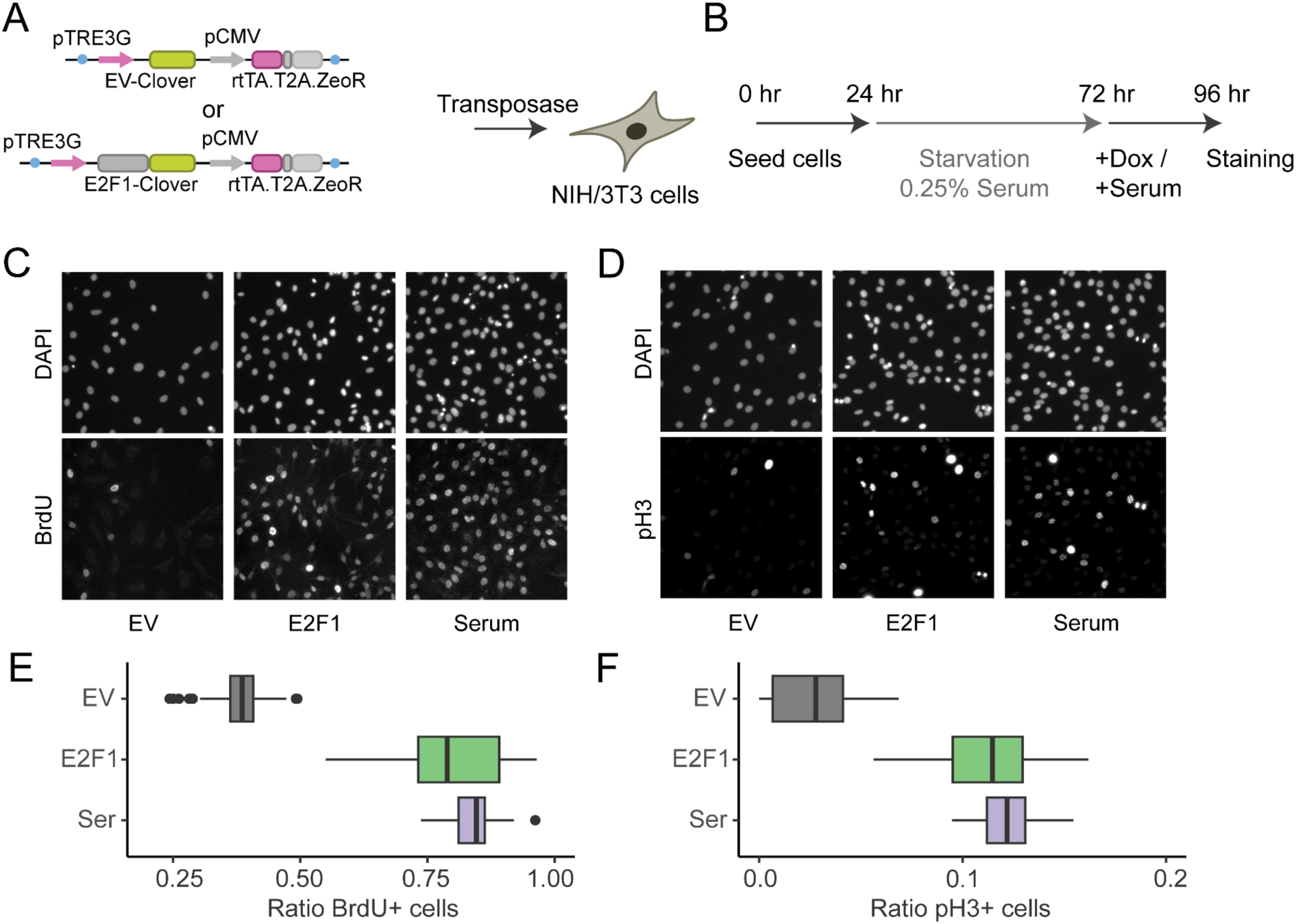
cell-cycle reentry by induced expression of E2F1. **A)** Plasmids for generation of NIH/3T3 cells with doxycycline-inducible expression of E2F1-Clover or negative Empty (EV) Vector control. **B)** Serum-starvation procedure to induce cell-cycle exit and measure reentry. **C)** Example DAPI and BrdU staining to measure S-phase entry in EV control induced, E2F1 induced, or 10% Serum treated cells. **D)** Example DAPI and pH3 staining to measure M-phase entry in EV control induced, E2F1 induced, or 10% Serum treated cells. **E)** Quantification of S-phase entry via BrdU staining, n = 3 independent experiments with 12-16 images each. **F)** Quantification of M-phase entry via pH3 staining, n = 3 independent experiments with 12-16 images each.

In addition to S-phase entry, E2F1 expression resulted in entry into G2/M phases of the cell-cycle as indicated by positive labeling for mitotic marker phospho-histone H3 (pH3) (**Figure 1D**). Quantification of our immunostaining showed that cells entered S-phase and G2/M phase with similar rates after E2F1 induction or serum addition (**Figure 1D-E**). Together, these results show that E2F1 expression is sufficient to trigger exit from quiescence and promote cell-cycle progression through G1, S, and G2 phases, leading to at least one complete round of cell division.

### E2F1 expression leads to rapid cell-cycle reentry

To monitor temporal dynamics of cell-cycle reentry at the single-cell level, we generated a dual reporter line carrying FUCCI-CDT1-mCherry to monitor cell-cycle progression and E2F1-Clover to measure E2F1 levels during cell-cycle reentry (**Figure 2A**). The FUCCI-CDT1-mCherry reporter is a fusion protein that is composed of a fragment of the DNA licensing factor (CDT1) fused to mCherry fluorescent protein, which undergoes degradation by SCF/Skp2 ubiquitin ligase as cells reenter the cell-cycle, resulting in loss of mCherry signal^26,27^. Using live single-cell fluorescent microscopy, we were able to associate levels of E2F1-Clover and simultaneously monitor cell-cycle reentry by measuring the reduction of mCherry signal at the single-cell level (**Figure 2B**). Using semi-automated image processing (**Supplementary Figure 2A**), we quantified the level of E2F1-Clover as well as FUCCI-CDT1 in individual cells^28^. We did not observe a noticeable decrease in the mCherry signal in the empty-vector (EV) control group, indicating that cells remained in quiescence (**Figure 2C, Movie 1**). For the E2F1 expressing cell line, the Clover measurements indicated that E2F1 levels peak approximately 5 hours after addition of dox, coinciding with cells reentering the cell-cycle as evidenced by a rapid and sharp decrease in mCherry signal (**Figure 2D, Movie 2**). While cells treated by 10% serum re-addition also showed a decrease in mCherry signal, they responded with several hours of delay when compared to the E2F1 expressing cells **(Figure 2E, Movie 3)**. No major differences in cell death were observed comparing induction of EV control, E2F1, or 10% serum treated cells, as measured by live cell propidium iodide imaging (**Supplementary Figure 2B)**. Like E2F1, induction of a closely related transcription factor in the activator E2F family, E2F2, resulted in cell-cycle entry from quiescence as measured by BrdU incorporation and pH3 labeling (**Supplementary Figure 1E-F)** and in live-cell FUCCI-CDT1-mCherry tracking **(Supplementary Figure 2C, Movie 4)**. Furthermore, the deletion of the DNA binding domain (DBD) of E2F1 led to a complete loss of E2F1 cell-cycle function as measured by BrdU, pH3 and the FUCCI-CDT1-mCherry sensor (**Supplementary Figure 1E-F, Supplementary Figure 2D, Movie 5)**, illustrating that DNA-binding is essential for transcriptional regulation of E2F1 target genes in the context of cell-cycle reentry. Therefore, we sought to use genome-wide transcriptome analysis to determine the E2F1-induced transcriptional programs that trigger cell-cycle reentry.

**Figure 2.**
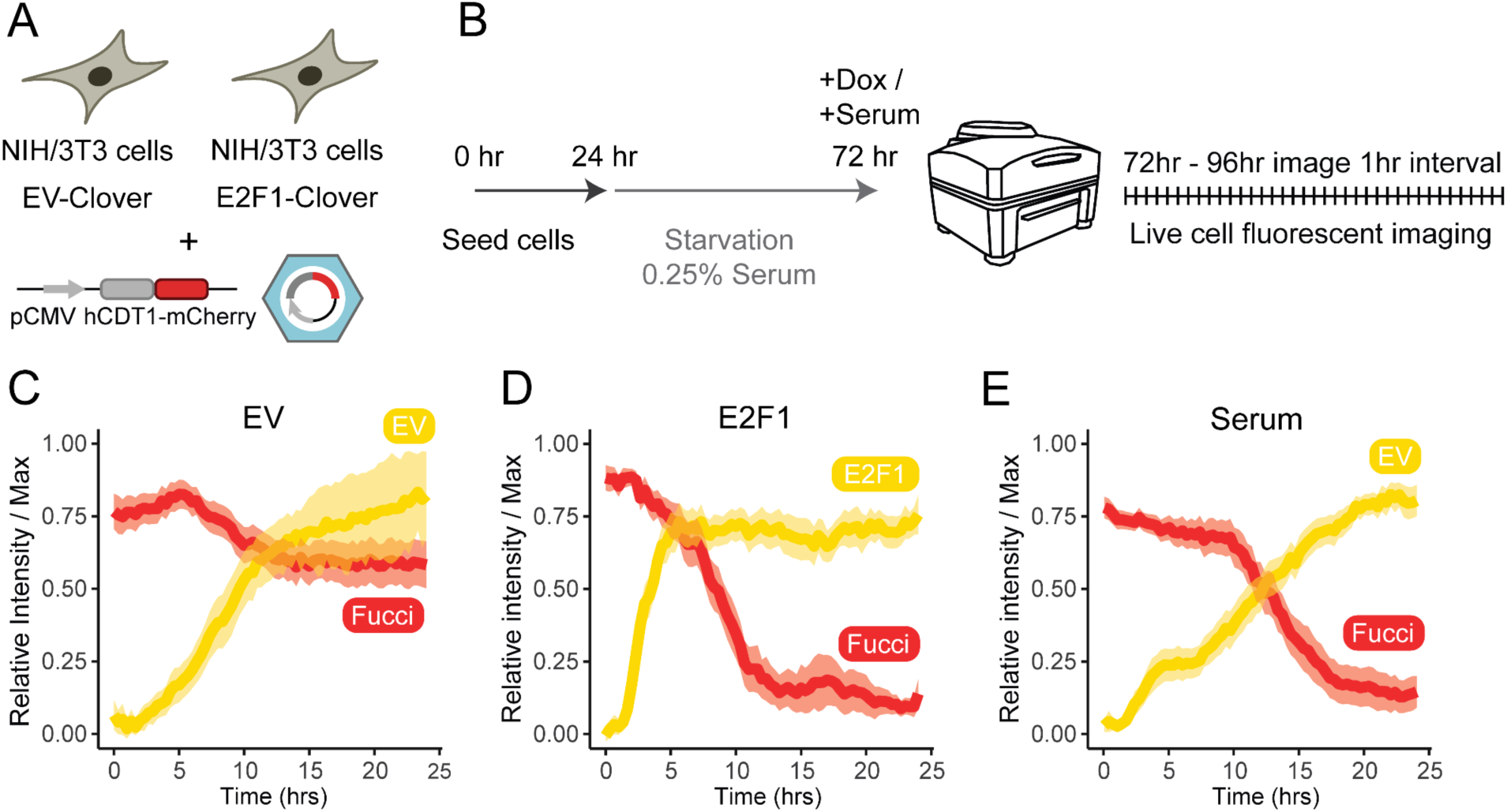
Dynamics of E2F1-mediated cell-cycle reentry. **A)** EV-Clover and E2F1-Clover lines transduced with CDT1-mCherry encoding lentivirus. **B)** Live cell imaging for dual capture of Clover and mCherry signals in double transgenic cell lines. **C)** Single-cell quantification of dox-treated EV-Clover::CDT1-mCherry cells (mean, 95% c.i., n = 73 individual cells). **D)** Single-cell quantification of dox-treated E2F1-Clover:CDT1-mCherry cells (mean, 95% c.i., n = 73 individual cells). **E)** Single-cell quantification of dox+serum-treated EV-Clover::CDT1-mCherry cells (mean, 95% c.i., n = 52 individual cells).

### Accelerated transcriptional programs of E2F1-induced cell-cycle reentry

To determine dynamics of the transcriptome as cells exit from the quiescent state to the G1-S-phase, we performed a time-course RNA-sequencing experiment at 0, 2, 6 and 12 hours after E2F1 induction. We also performed the same time-course RNA-sequencing experiment following 10% serum treatment, to compare mechanisms of cell-cycle reentry upon growth factor stimulation to that of E2F1 induction **(Figure 3A**). For each condition, we measured the relative expression and identified differentially expressed genes in comparison to the 0-hour time point. In the control condition (EV), only a handful of genes showed transcriptional changes, indicating that cells are in a relatively stable transcriptional state after 48 hours of serum starvation (**Figure 3B**). No significant associations were found using GO term analysis after k-means clustering for this small number of differentially regulated genes (**Supplementary Figure 3A**).

**Figure 3.**
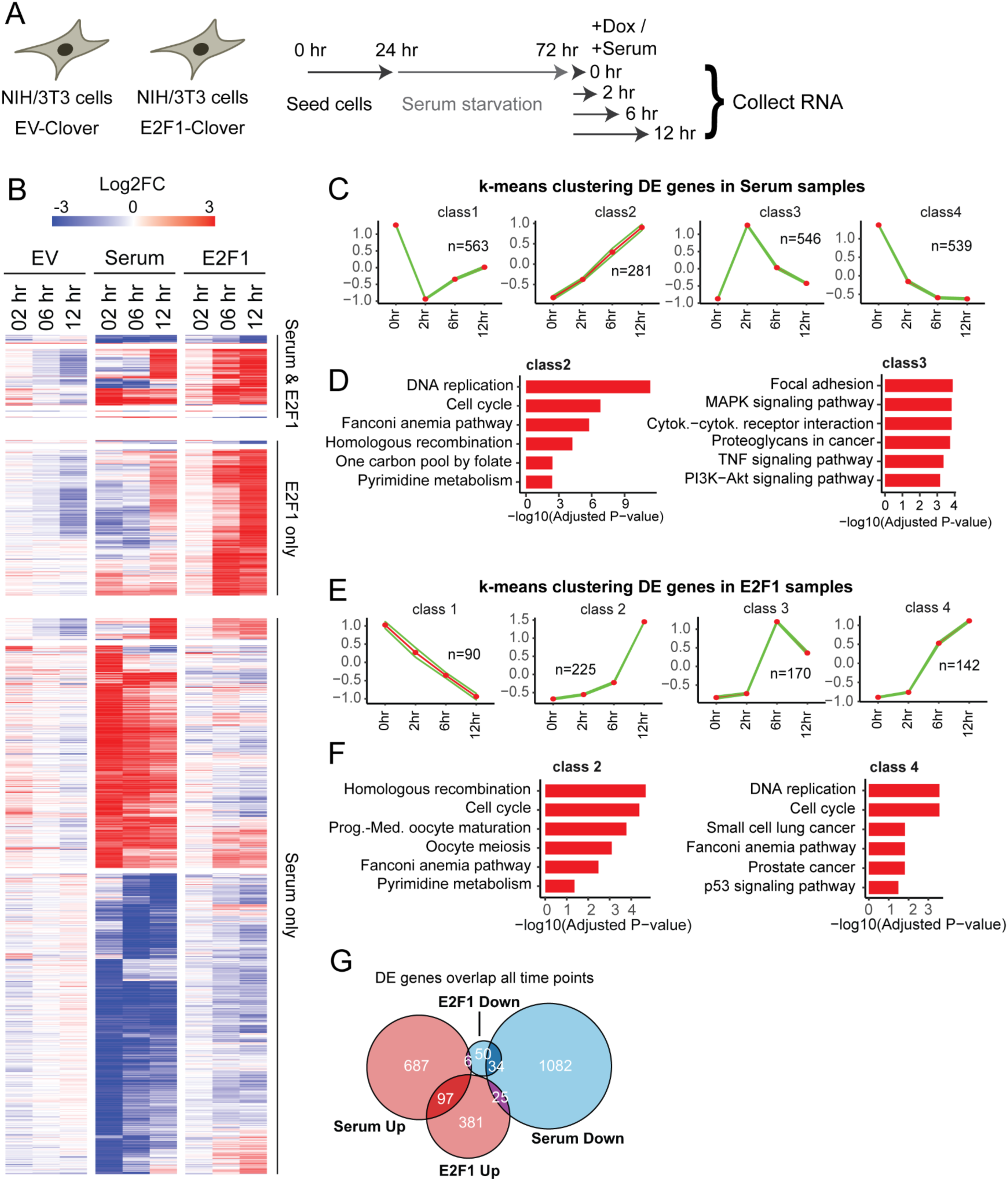
Transcriptome dynamics during exit from G0 by serum and E2F1. **A)** Experimental design for time-course RNA sampling. **B)** Heatmap showing differentially expressed genes (log2fc > 2, padj > 0.05) in Serum and/or E2F1 samples. **C)** K-means clustering of serum differentially expressed genes. **D)** Examples of Cluster 2 (slow regulated) and 3 (fast regulated) showing significant enrichment in GO term analysis. **E)** K-means clustering of E2F1 differentially expressed genes. **F)** Examples of Cluster 2 and 4 enrichment in GO term analysis related to cell-cycle processes. **G)** A Venn diagram of overlapping and distinct sets of genes between serum-induced and E2F1-induced cells.

In response to serum treatment, cells exhibit widespread transcriptional dynamics, including gradual upregulation, transient activation, and gradual decline in gene expression (**Figure 3C**). Hundreds of genes were up- or downregulated in roughly equal numbers within 2 hours of serum treatment, with many involved in growth signaling and homeostasis, while cell-cycle-related genes increased their activity at later time points (**Figure 3C-D**). In contrast, E2F1 induction mainly upregulates genes (**Figure 3E**), which is consistent with E2F1 acting as a transcriptional activator^23,29–31^. A small subset of genes was activated within 2 hours of E2F1 induction, this early response includes a 2-fold upregulation of canonical cell-cycle activators like Ccne1 (Cyclin E1) and Ccne2 (Cyclin E2), as well as other targets linked to the cell-cycle like the transcription factor Grhl1^32^ and the kinase Pim1^33^. The timing and identity of these genes suggests that they are likely key E2F1 direct target genes involved in the early response leading to cell-cycle reentry. More substantial transcriptional changes are observed following 6 and 12 hours of E2F1 induction (**Figure 2C**), mostly including genes associated with cell-cycle progression (**Figure 3E-F**).

Surprisingly, we found that there is relatively little overlap in E2F1 and serum differentially expressed genes (**Figure 3G, Supplementary Table 1**). Similarly, principal component analysis (PCA) showed that E2F-induced cells and serum-induced cells form distinct transcriptional clusters (**Supplementary Figure 3B)**. In part this result can be explained by the broader range of signaling pathways and subsequent activation of a global gene expression program in response to serum, compared to the more precise and targeted gene regulation using E2F1 only. Additionally, a subset of these genes could be induced at a faster rate in the E2F1-induced cells compared to serum treated cells. Our live imaging experiments showed that E2F1 induction resulted in accelerated cell-cycle reentry compared to serum treatment (**Figure 2C-E**). To investigate if cell-cycle progression is also accelerated on the transcriptome level, we analyzed cell-cycle associated genes. We found that canonical markers of cell-cycle progression, such as cyclins, show faster upregulation after E2F1 induction (**Supplementary Figure 3C**). Similarly, when analyzing a broader set of cell-cycle genes active during the G1/S and G2/M phases^34^, we observed that E2F1 induces their expression more rapidly than serum stimulation (**Supplementary Figure 3D**). Transcriptomic comparison shows that 12-hour E2F1-induced cells are only modestly correlated with 12-hour serum-treated cells. However, 6-hour E2F1 induction shows a much stronger correlation with 12-hour serum treatment (**Supplementary Figure 3E**), indicating that E2F1 triggers an accelerated cell-cycle transcriptional program. Together, these findings show that E2F1 not only activates a gene expression program partially distinct from that of serum stimulation but also does so with faster dynamics, promoting rapid progression through the cell cycle.

### E2F1 mediated cell-cycle reentry is not associated with global changes in chromatin accessibility

We were interested to see how the chromatin accessibility landscape is regulated in serum-starved cells after E2F1-induction or 10% serum treatment and how chromatin changes correspond with transcriptional activity of cell-cycle genes. Previous studies show that in quiescent cells, many cell-cycle genes are repressed to ensure fate stability and reentry into the cell-cycle involves re-activation of these genes^4,8,13,19,35^. However, it is not fully understood how changes in chromatin accessibility contribute to this transition. Specifically, it is not well understood whether chromatin must first become more accessible to enable transcription factor binding, or whether transcription factors can initiate gene activation without changing chromatin accessibility globally. Our gene expression analysis shows that E2F1-responsive genes, including many involved in cell-cycle progression, are rapidly upregulated upon E2F1 induction within 2-6 hours. Since gene activation is often assumed to depend on increased physical accessibility at promoters and distal elements^36^, we set out to examine whether E2F1 induces changes in chromatin accessibility as cells exit quiescence. To test this, we performed a time-course ATAC-seq experiment in parallel with the RNA-seq experiment, comparing chromatin accessibility dynamics in serum-treated cells, E2F1-induced cells, and empty vector (EV) controls.

First, we examined chromatin accessibility changes in promoters of genes that were differentially expressed. While serum treated cells globally increase chromatin accessibility for differentially expressed genes identified from RNA-Seq data in a time-dependent manner, E2F1 induction did not lead to global chromatin accessibility changes in differentially expressed genes compared to EV control (**Figure 4A, Supplementary Figure 4A**). Similarly, cell-cycle related genes did not show widespread differences in chromatin accessibility after E2F1 induction, but did generally show increased accessibility after serum treatment (**Supplementary Figure 4B-C**). Differential peak analysis of serum treated cells showed widespread changes in chromatin accessibility with thousands of regions gaining accessibility, which was not the case for E2F1 induced cells (**Figure 4B**). Motif analysis in serum-responding peaks showed they are most strongly associated with bZIP transcription factors, which are canonical downstream effectors for multiple growth factor signaling pathways^37^ (**Supplementary Figure 4D**).

**Figure 4.**
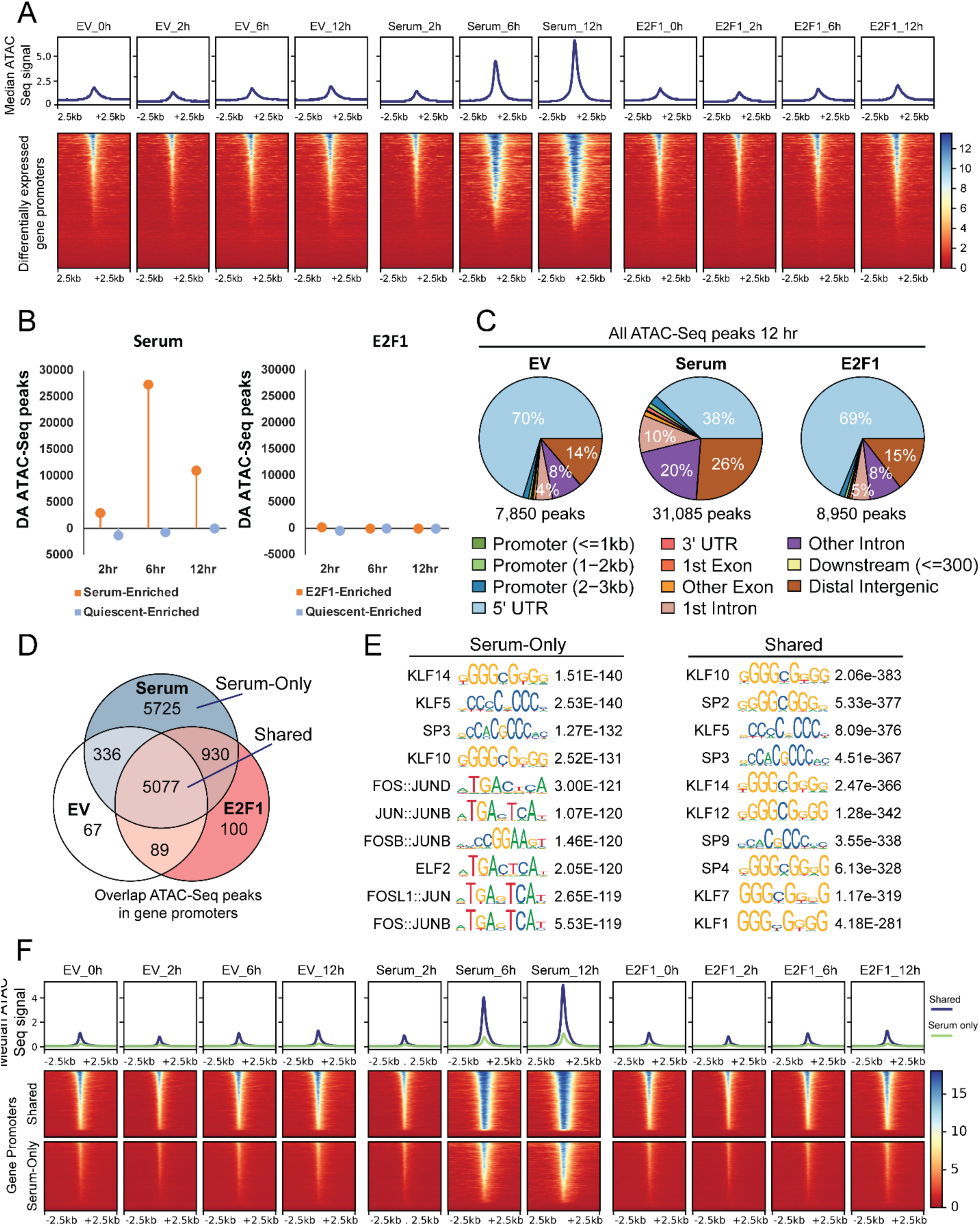
Dynamics of chromatin accessibility changes during exit from G0 by serum and E2F1. **A)** Chromatin accessibility in promoters of differentially expressed genes. **B)** Differential peak analysis relative to 0 hour data. **C)** Distribution of DNA annotations of all accessible regions at the 12 h time point in EV (left), serum treated (middle) and E2F1 induced (right) samples. **D)** Overlap in accessible gene promoters between conditions. **E)** Motif analysis showing top 10 motifs for Serum-only gene promoter accessible regions (left) and Shared gene promoter accessible regions (right). **F)** Profile and heatmap of ATAC-Seq data of shared promoter ATAC-Seq peaks (overlapping in EV, E2F1, serum), and serum-only promoter ATAC-Seq peaks.

Although E2F1 induction did not lead to widespread changes in chromatin accessibility, we detected low-level pre-existing ATAC-seq signals at a subset of promoter regions in G0 cells prior to E2F1 activation. We hypothesize that E2F1 can operate in a chromatin context with limited accessibility. To test this hypothesis, we first analyzed all accessible sites at the 12-hour time point and found approximately 8,000 accessible regions in both the EV and E2F1 conditions— most of which (∼70%) were promoter-associated compared to a nearly fourfold increase (31,000 regions) following serum stimulation, showing global reopening of proximal and distal regions (**Figure 4C**). Promoters that retain limited accessibility in quiescent cells showed strong overlap between EV and E2F1, whereas serum treatment led to the opening of an additional ∼5000 promoter regions (**Figure 4D**). Motif analysis in serum-only or shared accessible promoter regions (**Figure 4E, Supplementary Table 2**) revealed that, similar to previous motif analysis, serum-only peaks are enriched for bZIP factors in Fos-Jun complexes, as well as Ets factors like Elf2 that can act as pioneer factors^38^. Top motifs in the shared accessible promoters are mostly enriched for Sp/Klf sequences. Interestingly, proximal promoter regions with shared accessibility across conditions retain partial accessibility during quiescence in E2F-induced and EV control, whereas serum treated cells show much higher accessibility for all these regions (**Figure 4F).** These data suggest that E2F1 can operate in a permissive chromatin context with limited accessibility in quiescent cells and no widespread changes in chromatin accessibility are needed for E2F1 to fully commit cells to progress through the cell-cycle.

### E2F1 is able to directly interact with its binding sites located in regions with low-level accessibility

The lack of any obvious widespread E2F1-mediated changes in chromatin accessibility prompted us to test where E2F1 is able to access its binding sites near genes that show differential expression but have low-level accessibility signals and do not increase their chromatin accessibility. To measure E2F1 association at such sites, we used ChIP-Seq to map E2F1 binding sites after 12 hours of E2F1 induction in quiescent cells. Genome-wide analysis of E2F1 binding revealed that E2F1 directly accesses thousands of DNA regions (**Figure 5A**), predominantly at the promoters of genes it upregulates (80% bound), with minimal binding observed at promoters of downregulated genes (13% bound) based on transcriptome data (**Figure 5B**). Next, we found that serum treatment primarily induces chromatin accessibility at genes that are transcriptionally upregulated, with 75% of upregulated genes exhibiting increased accessibility, compared to only 20% of downregulated genes (**Figure 5C**). This result also suggests that a large fraction of genes upregulated by serum (∼25%) do not rely on substantial chromatin remodeling for activation, paralleling the E2F1-mediated gene activation, which occurs without widespread changes in chromatin accessibility.

**Figure 5.**
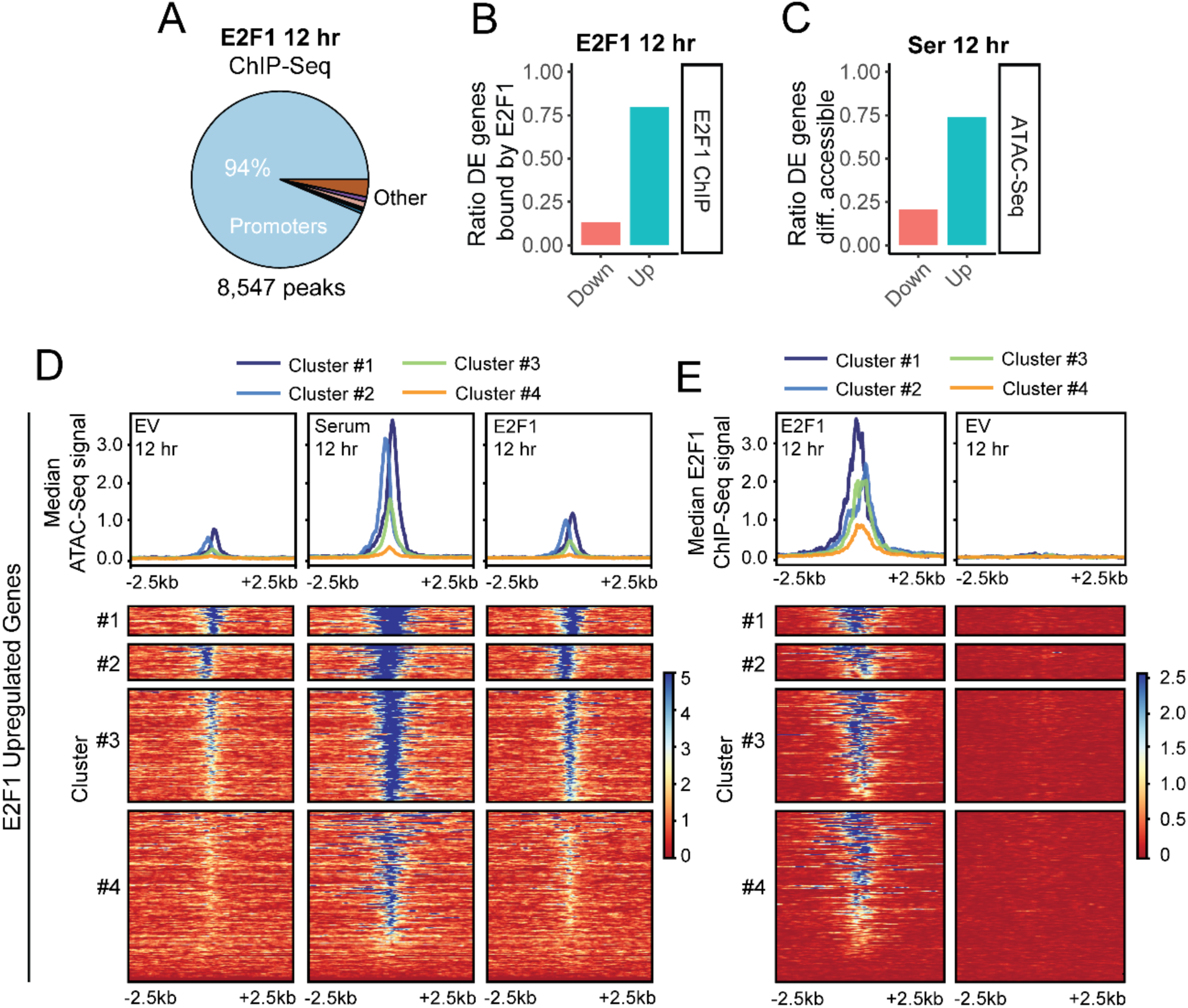
Genome-wide binding profile of ectopically expressed E2F1. **A)** Chart showing the distribution of E2F1 bound DNA annotation. **B)** Ratio of differentially expressed genes after E2F1 induction bound by E2F1. **C)** Ratio of differentially expressed genes after 10% serum treatment of E2F1 induction showing differential accessibility in serum data. **D)** Clustering of ATAC-Seq profiles in promoters of E2F1-upregulated genes, showing data of EV (left), 10% serum treated (middle) and E2F1 induced cells (right). **E)** E2F1 ChIP-Seq data mapped in the clustered promoter regions.

To look more closely at E2F1 upregulated genes, we clustered their promoter accessibility profiles into 4 groups that showed decreasing levels of accessibility (**Figure 5D**). Cluster 1 and Cluster 2 showed a detectable ATAC-Seq signal across conditions, whereas Cluster 3 appeared much lower in EV and E2F1 conditions and Cluster 4 accessibility was barely detectable for EV and E2F1 groups. Consistently, we found that nearly all promoters in cluster 1 and 2 show high E2F1-binding, but lower E2F1 binding signal in cluster 3 and 4 (**Figure 5E**). Interestingly, even though accessibility in cluster 4 E2F1 samples is very low, we still observe a relatively high amount of E2F1 binding signals in this cluster. These observations support the idea that low-level chromatin accessibility present during quiescence is sufficient for E2F1 binding and gene transcription, as exemplified by binding to the promoters of cell-cycle genes such as Pcna and Aurora Kinase B (Aurkb) and their gene expression upregulation (**Supplementary Figure 5, Supplementary Table 3**). Moreover, we identified regulatory elements of cell-cycle genes, such as Bub1, with a clear E2F1 binding peak, but no detectable ATAC-Seq peak signal (**Supplementary Figure 5, Supplementary Table 3).** Together, these findings show that E2F1 is able to bind and activate key cell-cycle genes with limited, or even no, chromatin accessibly. Surprisingly, increased chromatin accessibility is not broadly required for E2F1 function in gene activation and inducing cell-cycle entry. Moreover, E2F1-induced cells can complete a full cell-cycle without major reopening of chromatin, suggesting the cell-cycle might be uncoupled from other cellular homeostatic processes under our experimental conditions.

### E2F1 directly interacts with nucleosomes

It is thought that closed chromatin generally poses a barrier to transcription activation, as transcription factor binding sites are made inaccessible due to competition from nucleosomes^38^. However, recent examples demonstrate that some transcription factors can access DNA-binding motifs in the context of nucleosomes^39^. To test whether E2F1 is able to bind nucleosomes, we purified the E2F1 and DP2 DNA-binding domains (DBDs), which form a heterodimer, and assayed association with reconstituted nucleosomes that contain recombinant histones and the Widom 601 strong positioning DNA sequence with a TAMRA label (**Figure 6A**). Using a fluorescence polarization assay, we found that E2F1-DP2 DBDs bind weakly to the free DNA (K_d_ = 1.5 ± 0.1 μM) and form a tighter complex with the labeled nucleosome (K_d_ = 0.12 ± 0.01 μM), whereas the E2F1 transactivation domain (TAD), used as a negative control, shows no binding signal. While the 601 sequence does not contain a complete high-affinity E2F consensus site (TTTCGCGCG), it does contain sequences that nearly match the minimal E2F binding element (CGCGCG), perhaps explaining the observed weak affinity association with the free Widom 601 DNA. The notable increase in affinity upon binding the reconstituted nucleosomes suggests that additional interactions with the histone core are present.

**Figure 6:**
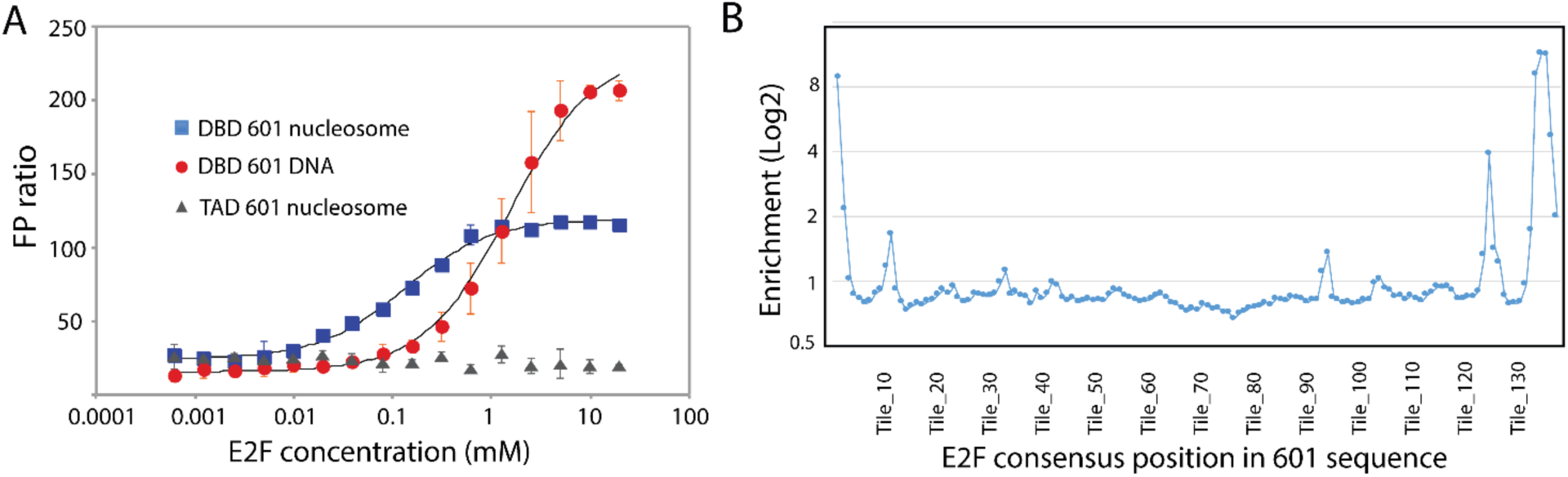
E2F binding to nucleosomes. **A)** Fluorescence polarization (FP) titration of E2F1-DP2 DNA binding domains (DBDs) or E2F1 transactivation domain (TAD) control into FITC-labeled 601 DNA or 601 nucleosomes. Error bars are standard deviations from three experiments. **B)** Nucleosome binding preferences of E2F1-DP2 DBDs determined by SeEN-seq. The plot displays the relative enrichment of E2F1-DP2 binding to nucleosomes containing the E2F consensus motif (mean of 3 independent experiments), systematically tiled across the 601-nucleosome positioning sequence. The X-axis indicates the 601 base pair position at which the first base of the consensus motif is located.

In order to observe whether the E2F consensus motif can be recognized in the context of a nucleosome and whether there is a preference for location, we used the high-throughput sequencing approach called Selected Engagement on Nucleosome sequencing (SeEN-seq)^40^. E2F1-DP2 DBD was incubated with a library of nucleosomes in which the E2F consensus motif was tiled through the Widom 601 sequence. Following separation of bound and unbound species by native PAGE electrophoresis, extracted DNA was subjected to next-generation sequencing (NGS), allowing for the calculation of relative enrichments of E2F1-DP2 binding at each motif position throughout the nucleosome (**Figure 6B**). As previously observed for several other transcription factors^40–43^, the data demonstrate a preference for motif positions near the ends of the nucleosome wrap. Remarkably, we also observe a periodic preference for sites that are spaced ∼12 nucleotides apart and that occur symmetrically from the ends toward the center of the nucleosome. This pattern is consistent with a preference for binding motif positions with a specific orientation on nucleosomal DNA. Together, biochemical data support a mechanism in which E2F1 can bind to consensus sites in the genome even in the presence of nucleosomes.

## Discussion

Many cells in multicellular organisms reside in a relatively stable non-dividing quiescent state of the cell-cycle for most of their life. While quiescent cells are transcriptionally active, they repress genes that are required for cell-cycle progression through multiple epigenetic layers that are not fully understood. It is commonly assumed that activation of these genes requires global re-organization of chromatin accessibility, followed by transcription factor binding and recruitment of the transcription machinery.

Indeed, when we used growth factor stimulation (serum addition) to trigger cell-cycle reentry, we observed such global changes in chromatin accessibility. However, quiescent cells can reenter the cell-cycle and complete their division without global changes in chromatin accessibility when induced by expression of a single transcription factor; E2F1. E2F1 induced cell-cycle reentry is faster than growth factor (serum) induced cell-cycle reentry and associated with a transcriptional program that displays distinct temporal dynamics and features compared to the program induced by serum. Surprisingly, E2F1 is able to activate gene expression programs and elicit cell fate changes without changing global chromatin accessibility. E2F1 may take advantage of pre-existing low level accessibility in cell-cycle gene promoters to access binding sites. In fact, the DREAM complex, which contains the E2F family repressor E2F4, binds promoters, including cell cycle genes, when G1-S genes are repressed during quiescence, suggesting some degree of E2F-binding site accessibility^8,13,14^. However, we also identified promoters lacking a clear ATAC-Seq peak where E2F1 binding still occurs, consistent with the possibility that E2F1 can directly engage nucleosomes to access its sites. In line with this model, we found that E2F1 is able to bind and modify chromatin configuration through direct interaction with nucleosomes. The precise mechanisms underlying the transcriptional activation of cell-cycle genes by E2F1 without significant changes in chromatin accessibility remain unknown and merits further investigation. However, DNA binding is essential for this function, as deletion of the DNA binding domain in E2F1 led to a complete loss of cell-cycle reentry. One hypothesis is that local modification of nucleosome positioning by E2F1 is sufficient to activate gene expression without affecting global chromatin accessibility as measured by ATAC-Seq. In this sense, E2F1 may use a different mechanism to activate transcription compared to TFs that promote increased accessibility upon binding to regulatory regions.

We predict that there are other transcription factors that might trigger gene expression without changing chromatin accessibility as it has recently been shown that there is noticeable discordance between gene expression and chromatin accessibility^44^. The discordance between changes in chromatin accessibility and gene expression is also observed in our serum treated cells where we observed that 25% of upregulated genes show differential gene expression without changing their chromatin accessibility. This result suggests a subset of genes may be regulated independent of changes in local chromatin accessibility and through accessing their binding sites on nucleosomal DNA to activate gene expression from repressed regions of the chromatin. We speculate that such mechanisms are employed by transcription factors to drive cell fate changes during natural development or reprogramming to activate genes that are not accessible for transcription. While E2F1 shares key features with pioneer factors, such as the ability to bind nucleosomes and activate transcription from chromatin with limited accessibility to drive cell fate transitions, it differs in that, under our experimental conditions, it does not induce broad changes in chromatin accessibility, suggesting its pioneer-like activity may depend on specific cellular or chromatin contexts.

## Methods and Materials

### Cell Culture

NIH-3T3 cells were grown in DMEM supplemented with 10% heat-inactivated fetal bovine serum (HIFBS) and 1x penicillin+streptomycin (P/S). For maintaining cultures, cells were split using 0.25% Trypsin + 0.5mM EDTA every 3-4 days at a subcultivation ratio of 1/10.

To generate doxycycline-inducible stable cell lines, PiggyBac transposon integration was used. NIH/3T3 cells were seeded at a density of 200,000 cells per well in a 6 well plate. After 24 hours, medium was replaced with 2ml DMEM + 10% HIFBS. Then, transfection complexes were made containing the following for transfecting one well: In 1.5ml eppendorf tube 1: 100μl OptiMEM, 200ng PiggyBac transposase plasmid, 800ng PiggyBac transposon TRE3G-E2F1-Clover, 1000ng pUC19 plasmid. In 1.5ml eppendorf tube 2: 100μl OptiMEM + 8 μl TurboFect. Both tubes were briefly vortexed and centrifuged. 100ul of TurboFect mix from tube 2 was added into plasmid mix tube 1. The tube was briefly vortexed and centrifuged, incubated 20 min at room temperature, and the complete volume was added to one well of the 6 well plate. 6 hours after adding transfection complexes, 1ml DMEM + 10% HIFBS + 3x P/S was added. 48 hours after transfection, Zeocin was added to the culture medium at 400 μg/ml. Medium was changed every 2-3 days and supplemented with Zeocin, until control cells (cells transfected with PiggyBac transposon TRE3G-E2F1-Clover only) were dead.

For experiments with serum starvation, 30,000 NIH/3T3 cells were seeded per well in a 24 well plate in DMEM + 10% HIFBS + 10x P/S. 24 hours after seeding, medium was replaced with 550 ul starvation medium (DMEM + 0.25% HIFBS + 1x P/S). 48 hours after adding starvation medium, one of the following was added per well as indicated: 50 μl of starvation medium (-Dox), 50 μl of starvation medium + 5000 ng/ml doxycycline (+Dox), or 50 μl HIFBS (+Serum). For BrdU labeling experiments, these were supplemented with 10mM BrdU. Final concentrations were approximately 500 ng/ml doxycycline, 10% HIFBS, and 1mM BrdU after addition into the wells already containing starvation medium. Under these conditions, cells were cultured for up to 24 hours before isolation.

### Live Imaging

To measure cell-cycle in live cells, the FUCCI-hCDT1-mCherry sensor was integrated via lentivirus transduction into the previously used doxycycline-inducible E2F1-Clover cell lines.

Lentivirus was made using the following protocol: HEK293T cells were seeded at a density of 500,000 cells per well in a 6 well plate, in DMEM + 10% FBS + 1x P/S. After 24 hours, medium was replaced with 2ml DMEM + 10% HIFBS. Then, transfection complexes were made containing the following for transfecting one well: In 1.5ml eppendorf tube 1: 100ul OptiMEM, 300ng VSVG plasmid, 450ng DR8.74 and 750ng lenti-FUCCI-hCDT1-mCherry plasmid. In 1.5ml eppendorf tube 2: 100μl OptiMEM + 8 μl TurboFect. Both tubes were briefly vortexed and centrifuged. 100μl of TurboFect mix from tube 2 was added into plasmid mix tube 1. The tube was briefly vortexed and centrifuged, incubated 20 min at room temperature, and the complete volume was added to one well of the 6 well plate. 6 hours after adding transfection complexes, 1ml DMEM + 10% HIFBS + 3x P/S was added. 72 hours after transfection, culture medium was collected in a 15ml tube and centrifuged for 5 min at 500 rcf. Medium supernatant was filtered using a 0.45μm syringe filter into a new tube and stored at 4°C. To transduce NIH/3T3 cells, they were seeded at a density of 200,000 cells per well in a 6 well plate. After 24 hours, the medium was replaced with 1.5ml DMEM + 10% HIFBS + 1x P/S, and 1.5ml of filtered lentivirus medium. Cells were incubated for 24 hours with lentivirus, after which the medium was replaced with DMEM + 10% HIFBS + 1x P/S again.

For live imaging of cells with FUCCI-hCDT1-mCherry sensor, cells were grown in 24 well plates and underwent the starvation procedure as described before. Upon addition of doxycycline, serum or control solutions, cells were immediately transferred to a live cell imaging microscope (BioTek Lionheart FX), using the humidity incubator chamber at 37°C (2°C gradient) and 5% CO2, and imaged every 20 minutes.

For live imaging of cells with propidium iodide staining, cells were grown in 24 well plates and underwent the starvation procedure as described before. Upon addition of doxycycline, serum or control solutions, propidium iodide was also added to 1 ug/ml. Cells were immediately transferred to a live cell imaging microscope (BioTek Lionheart FX), using the humidity incubator chamber at 37°C (2°C gradient) and 5% CO2, and imaged every 30 minutes.

### ATAC-Seq, RNA-Seq, ChIP-Seq experimental workflow

NIH/3T3 wells were cultured as described before using the 48 hour serum starvation procedure. Cells were collected at the end of serum starvation (t=0), and at 2, 6 and 12 hours after adding doxycycline or serum. Cells for ATAC-Seq and RNA-seq were grown in parallel, starting with 2 wells each in a 6 well plate for each timepoint, with an initial seeding density of 60,000 cells per well. Replicates one and two were collected from 2 independent experiments following the same protocol.

For ATAC-Seq, at each timepoint, medium was removed from wells and washed once with 1x PBS. Cells were detached using 1ml 0.25% Trypsin + 0.5mM EDTA, after which 1ml of DMEM + 1% HIFBS + 1x P/S was added. Cells were centrifuged and resuspended in 1ml DMEM + 0.25% HIFBS + 1x P/S. Cell number and viability was measured using an automated cell counter (Bio-Rad TC20). 100,000 live cells were collected in a 1.5ml microcentrifuge tube and processed for ATAC-Seq using the provided protocol (Active motif 53150). Prepared ATAC-Seq Libraries were analyzed using a Bioanalyzer 2100 (Agilent), using the DNA high sensitivity chip and reagents (Agilent 5067-4626). Samples were sequenced using the Illumina NovaSeq 6000 platform, paired-end with 150bp reads (Novogene).

For RNA-Seq, at each timepoint, medium was removed from wells, washed once with 1x PBS, followed by addition of 400ul of RNA lysis buffer (Zymo R1051) per well and incubated 2 minutes at room temperature. Lysed cells were transferred to a 2ml eppendorf tube, the lysate from the duplicate wells were combined into the same tube. RNA was extracted using Zymo Quick RNA microprep spin column purification including DNAseI treatment (Zymo R1051). RNA-Seq library preparation (PolyA selection, unstranded) and sequencing (150bp paired-end, Illumina NovaSeq 6000) was done by Novogene.

### Computational analysis of next generation sequencing data

RNAseq FASTQ files were inspected with FastQC. Low quality reads were removed using Trimmomatic, removing adapter/index sequences, quality filtering (sliding window 4:20) and retaining only reads of 50bp or longer. Data was aligned using HISAT2, unstranded, to the mouse mm10 genome. Featurecounts was used for read counts per gene, with Gencode VM23 gene annotation. Count normalization and differential gene expression was calculated using DESeq2. Differential genes per condition were subjected to k-means clustering into 4 groups, followed by gene-ontology analysis.

ATAC-seq FASTQ files were inspected with FastQC. Low quality reads were removed using Trimmomatic, removing adapter/index sequences, quality filtering (sliding window 4:20) and retaining only reads of 50bp or longer. Data was aligned using Bowtie2, to the mouse mm10 genome. Reads flagged as PCR duplicates were removed using RmDup, and reads in ENCODE blacklisted regions were removed (mm10.blacklist.v2). MACS2 was used for peak calling, peaks with q value > 50 were used for later analysis. Pooled replicate MACS2 analysis used to combine data from biological replicate experiments. HOMER was used for differential peak signal and motif calling. ChIPSeeker was used to classify peaks based on Gencode VM23 gene annotation. For visualization, gene promoters of differentially expressed genes were extracted by retrieving intervals 2.5kb upstream and downstream of the transcription start sites, followed by plotting the ATAC-Seq signals in these intervals using deepTools.

ChIP-seq FASTQ files were inspected with FastQC. Low quality reads were removed using Trimmomatic, removing adapter/index sequences, quality filtering (sliding window 4:20) and retaining only reads of 50bp or longer. Data was aligned using Bowtie2, to the mouse mm10 genome. MACS2 was used for peak calling, and only peaks with a q-value > 13 were used for analysis. ChIPSeeker was used to classify peaks based on Gencode VM23 gene annotation. To correlate chromatin accessibility (ATAC-Seq) with E2F1 binding (ChIP-Seq), differentially expressed genes in doxycycline-induced E2F1 conditions were analyzed. The gene promoters of this set were extracted, and the ATAC-Seq signal was mapped and clustered using deepTools, via k-means clustering into 4 groups. The E2F1 ChIP-Seq signal was then mapped with deepTools according to this clustering.

### Immunostaining

NIH/3T3 cells were grown in 24 well plates as described before. Culture medium was removed and wells were washed once with 1x PBS. Per well, 400ul of 3.6% formaldehyde in 1x PBS was added and incubated for 10 min at room temperature. Formaldehyde solution was removed and cells were washed twice with 1x PBS. Then, cells were permeabilized by incubation for 20 min in 1x PBS + 0.2% Triton. For BrdU staining, permeabilization solution was removed, cells were washed once in 1M HCl solution, followed by adding 400ul of 1M HCl solution and incubated for 40 min at room temperature. Then, HCl solution was removed, cells were washed once with 1x PBS, followed by addition of 400μl blocking solution containing Bovine Serum Albumine (BSA) per well (1x PBS + 3% BSA + 0.1% Triton) and incubated for 1 hour. Blocking solution was removed, 400ul antibody incubation solution was added per well (1x PBS + 1% BSA) and incubated overnight at 4°C. For other antibody stainings not requiring HCl treatment, the HCl wash and incubation was not done and well were incubated with blocking solution following permeabilization instead. Primary antibodies added into the antibody incubation solution were either mouse anti-BrdU (1:2,000) or mouse anti-pH3 (1:2,000). The next day, antibody solution was removed and wells were washed twice with 1x PBS + 0.1% tween (1x PBST) and once with 1x PBS. 400μl of antibody solution was added per well, containing 1:1,000 goat anti-mouse-alexa 594 secondary antibody and incubated for 1 hour at room temperature. Antibody solution was removed and washed once with 1x PBST. Then, 400μl of PBST + 0.5μg/ml DAPI was added per well and incubated for 5 min at RT. 1x PBST + DAPI solution was removed and wells were washed twice with 1x PBS, followed by addition of 500μl 1x PBS per well. Imaging of immunofluorescence stainings was done using the Biotek Lionheart FX.

### Protein expression and purification

The human E2F1 and DP2 DBDs were expressed in *Escherichia coli* from an engineered pGEX plasmid with an N-terminal GST tag and a TEV protease cleavage site. Protein was expressed overnight by inducing with 1 mM IPTG at 19 °C. All proteins were lysed in a buffer containing 500 mM NaCl, 40 mm Tris pH 8.0, 5 mM DTT, and 1 mM PMSF. The lysed cells were clarified by centrifugation at 19,000 rpm for 45 min at 4 °C. Protein lysates were allowed to bind to equilibrated glutathione sepharose resin (Cytiva) for 30 min and washed to remove unspecific proteins. The protein was eluted with a buffer containing 200 mM NaCl, 40 mM Tris pH 8.0, 5 mM DTT, and 10 mM reduced L-Glutathione. Eluted proteins were further purified using Q-sepharose and cleaved with TEV protease at 4 °C overnight. Proteins were then passed through glutathione sepharose resin to remove the free GST and concentrated to run through Superdex-75 (GE Healthcare) into 200 mM NaCl, 25 mM Tris pH 8.0, and 1 mM DTT. E2F1 and DP2 were mixed in a 1:3 molar ratio and incubated for 30 minutes on ice before isolating the E2F1-DP2 complex by passing through a Superdex-200 column into 200 mM NaCl, 25 mM Tris, and 1 mM DTT (pH 8.0). The Xenopus laevis H2A, H2B, H3, and H4 histones were expressed in *Escherichia coli* from a pET vector in pLysS cells. The cells were grown in LB media, induced at OD600 0.6-0.9, and induced with 0.4 mM IPTG. Cells were harvested after 3 hours of incubation at 37 °C. The bacterial pellets were resuspended in lysis buffer containing 1 M NaCl, 50 mm Tris pH 7.5, 1 mM BME, and 1 mM EDTA. Resuspended pallets were flash-frozen in liquid nitrogen and stored at -20°C overnight. Cells were lysed using a sonicator and centrifuged at 19,000 rpm for 30 minutes. Pellets containing inclusion bodies were washed with lysis buffer by mincing with a dounce homogenizer. This step was done 3-4 times with centrifuging at 19,000 rpm for 30 minutes. The pellet was then resuspended in a buffer containing 7 M guanidine hydrochloride and 20 mM Tris-HCl (pH 7.5) and incubated at 37 °C for 30 mins to extract histones. Undissolved material from the sample was removed by centrifugation and injected into a Sephacryl S200 column to separate histones from unspecific DNA. Fractions containing histones were dialyzed into a buffer containing 7 M freshly de-ionized urea, 20 mM sodium acetate pH 5.2, 200 mM NaCl, 1 mM EDTA, 5 mM BME, and 10 mM DTT. The protein was loaded onto a Source S cation exchange column and eluted as a gradient in the same buffer with 1 M NaCl. Fractions containing the histones were dialyzed against water with 1 mM BME for 72 hours with 3-4 buffer changes. Histones were then run on an SDS-PAGE to check for purity and lyophilized for storage. Histone octamer was reconstituted by mixing equimolar amounts of histone in a buffer containing 7 M guanidine hydrochloride, 20 mM Tris pH 8.0, and 10 mM BME. The undissolved histones were removed by centrifugation, and the supernatant was dialyzed in a buffer containing 20 mM Tris pH 8.0, 2 M NaCl, 1 mM EDTA and 10 mM BME for 72 hours with 3-4 buffer changes at 4°C. The octamer was then eluted from a Superdex 200 column in a buffer containing 20 mM Tris pH 7.5, 2 M NaCl, 1 mM EDTA and 10 mM BME. The octamer was stored at -80°C with 15% glycerol.

### Nucleosome reconstitution

The nucleosome core particle (NCP) was reconstituted using the Widom 601 positioning sequence. DNA was amplified by PCR with the forward primer 5’ ATCCCTATACGCGGCCGCCCTGGA-3’ and reverse primer (TAMRA)-5’ ACAGGATGTATATATCTGACACGTG-3’. The DNA was purified by gel-extraction. Following purification, the DNA was resuspended with equimolar amounts of histone octamer in a buffer containing 20 mM Tris pH 7.5, 2 M NaCl, 1 mM EDTA and 10 mM BME. The suspended octamer and DNA mixture was then dialyzed into the same buffer with decreasing NaCl concentration starting from 1 M, 0.75 M, 0.5 M, 0.3 M, 0.2 M, and finally 0.1 M with at least 6 hours of dialysis at 4°C. The NCP was then eluted from a Superdex 200 column and analyzed using 5% native PAGE. Nucleosomes were purified on a monoQ 5/50 ion exchange column, and nucleosome containing fractions were dialyzed into a buffer containing 20 mM Tris pH 7.5, 0.1 M NaCl, 1 mM EDTA, and 1 mM BME and stored with 15% glycerol at -80°C.

### Fluorescence Polarization (FP) assay

Dissociation constants for direct binding between E2F1-DP2 DBD and nucleosomes were determined by titrating increasing amounts of E2F1-DP2 DBD into 20 nM of TAMRA-labeled nucleosomes in a buffer containing 100 mM NaCl, 20 mM Tris pH 7.5, 1 mM DTT, and 0.1% Tween20. FP measurements were acquired on a PerkinElmer EnVision 2103 Multilabel plate reader with excitation at 559 nm and emission at 580 nm. The dissociation constants (K_D_) were calculated by fitting millipolarization (mP) values of three technical replicates against concentration using a one-site–binding model in GraphPad Prism 8. The Widom DNA probe was amplified by using a forward primer containing TAMRA dye and the following sequence: /TAMRA//iSp9/CTGGAGAATCCCGGTGCCGAGGCC (synthesized by Integrated DNA Technologies).

### SeEN-seq reagent preparation

The E2F consensus motif (TTTGGCGCC) was systematically tiled through the 147 bp Widom 601 nucleosome positioning sequence at single base-pair intervals, generating a library of 139 distinct DNA sequences with varying motif positions. These library sequences were flanked by EcoRV restriction sites and adapter sequences and synthesized as gene fragments by TWIST Biosciences. After resuspension, the individual fragments were pooled in equimolar amounts and digested with EcoRV-HF (NEB). The resulting 153 bp DNA fragments were purified from a 3% agarose gel using the QIAquick Gel Extraction Kit (Qiagen). For nucleosome reconstitution, E2F motif-containing DNA fragments were combined with an excess of unmodified Widom 601 DNA at a 1:30 molar ratio.

The DNA library was combined with assembled histone octamers at a 1:1.5 molar ratio in the presence of 2 M KCl. This mixture was placed in high-salt buffer (10 mM Tris-HCl pH 7.5, 2 M KCl, 1 mM EDTA, 1 mM DTT), and the salt concentration was gradually reduced to 0.25 M over three days at 4°C using a peristaltic pump and low-salt buffer (10 mM Tris-HCl pH 7.5, 250 mM KCl, 1 mM EDTA, 1 mM DTT). Following dialysis, the reconstituted nucleosomes were incubated at 55 °C for one hour. The nucleosome pool was then purified using a MonoQ 5/50 ion exchange column pre-equilibrated with buffer A (20 mM Tris-HCl pH 7.5, 100 mM NaCl, 0.5 mM TCEP) and eluted with a gradient increasing to 100% buffer B (20 mM Tris-HCl pH 7.5, 1 M NaCl, 0.5 mM TCEP). Fractions containing octameric nucleosomes were pooled, buffer-exchanged using a 30 kDa cut-off Amicon concentrator (Millipore) into 20 mM Tris-HCl pH 7.5 and 0.5 mM TCEP, and concentrated to 30 µl.

### SeEN-seq assay

SeEN-seq was performed as previously described^40^ with minor modifications. Nucleosomes (100 nM) were incubated with 2 µM E2F1-DP2 DBDs for 1 hour at room temperature in 20 µL reactions containing 20 mM Tris-HCl pH 7.5, 75 mM NaCl, 10 mM KCl, 1 mM MgCl₂, 0.1 mg/mL BSA, and 1 mM DTT. Three technical replicates were prepared. Samples were resolved on 6% native polyacrylamide gels (acrylamide:bis-acrylamide 37.5:1) in 0.5× TGE buffer for 1.2 hours at 150 V (room temperature). Gels were stained with SYBR Gold nucleic acid stain (10 min, Invitrogen), and DNA bands corresponding to TF-bound and unbound nucleosomes were excised. Gel slices were incubated in 100 µL acrylamide extraction buffer (500 mM ammonium acetate, 10 mM magnesium acetate, 1 mM EDTA, 0.1% SDS) for 30 minutes at 50 °C. Following this, 50 µL H₂O and 450 µL QG buffer (QIAquick Gel Extraction Kit, Qiagen) were added to each sample and incubated for another 30 minutes at 50 °C. After a short spin, the supernatant was transferred to QIAquick spin columns, and DNA was purified according to the manufacturer’s instructions. DNA was eluted in 22 µL H₂O, and approximately 10 µL (1–10 ng DNA) were used for NGS library preparation using the NEBNext Ultra DNA Library Prep Kit (E7370S/L) with dual indexing (E7600S) and 12 PCR amplification cycles. Purified libraries were quantified using a Qubit fluorometer (Thermo Fisher), pooled in equimolar ratios, and assessed for size distribution using a High Sensitivity DNA assay on an Agilent 2100 Bioanalyzer. Sequencing was performed on an Illumina NextSeq platform (300 bp paired-end). Sequencing reads were aligned to the reference DNA library using the Bioconductor package QuasR with default settings^45^, which employs Bowtie for read mapping^46^. Read counts for each construct were quantified using the Qcount function. To calculate SeEN-seq enrichment, library-size-normalized read counts were compared between TF-bound and unbound nucleosome fractions for each 601-E2F variant.

### Image Quantification

For quantification of end-point stainings (DAPI, BrdU and pH3), images were analyzed using imageJ (FIJI^47^). For BrdU / DAPI ratio, images from BrdU and DAPI labeling were segmented via thresholding, using a fixed threshold across conditions. BrdU images were subjected to background subtraction (50px radius) before thresholding. The number of particles was counted for each image, retaining only particles between 200-1200 pixel^2^. and the ratio of BrdU particles over DAPI positive particles was calculated. The pH3 / DAPI ratio was calculated using the same method. The speckled patterned nuclei of pH3 stained cells were merged into a single particle using the ImageJ “Dilate” option repeatedly on thresholded data, to avoid counting multiple particles per nucleus.

To segment and track single cells expressing the FUCCI reporter in live imaging experiments, we developed a MATLAB-based application **(Figure S2A)** that is specifically designed for the analysis of live imaging of cells expressing fluorescent FUCCI reporter along with other fluorescent markers. This application employs a hybrid automated and manual process for image segmentation and tracking. It initially separates different objects within the two-channel images and subsequently measures the intensity of the fluorescent signal in each channel. This MATLAB-based application streamlines FUCCI cell image analysis by enabling label assignment to each object for precise tracking and segmentation. To achieve accurate segmentation, this application harnesses various morphological image functions along with the watershed technique. Users can easily upload images, track cell attributes, and export quantification data in various formats. The codes and installable MATLAB application is available here: https://github.com/abzargar/FUCCI_Annotation_App.

### Statistical Analysis

## Data and Code availability

All raw and processed high-throughput sequencing data generated in this study have been deposited under the accession number: GSE296085 for RNA-Seq, GSE296087 for ChIP-Seq, GSE296086 for ATAC-Seq. All the time-lapse movies and microscopy images are available under Zenodo DOI: xxx. Our imaging annotation software is uploaded at: https://github.com/abzargar/FUCCI_Annotation_App. Any additional information required to reanalyze the data reported in this work paper is available from the Lead Contact upon request

## Resource availability

All unique/stable materials generated in this study are available from the lead contact upon reasonable request with a completed materials transfer agreement.

## Acknowledgement

We would like to thank all the current and former member of the Dr. Shariati’s and Dr. Rubin’s laboratories. This work was supported by the NIGMS/NIH through a Pathway to Independence Award K99GM126027/R00GM126027 (S.A.S.), NIGMS/NIH Maximizing Investigator Awards R35GM147395 (S.A.S.) and R35GM145255 (S.M.R), P01CA254867 (S.M.R.), and start-up package of the University of California, Santa Cruz (S.A.S), Gerrald Lodewijk is supported by a Dutch Research Council (NWO Domain Science Board) award (2021/ENW/01120083). J.W and N.H.T are supported by European Research Council (ERC) under the European Union’s Horizon 2020 research program (NucEM, No. 884331) and Swiss National Science Foundation SNF#: 310030_214852. Ben Topacio is supported by the NIH IRACDA fellowship (K12GM139185). We acknowledge core support from the UCSC Institute for the Biology of Stem Cells (IBSC), IBSC imaging facility (SCR_021135), CIRM Shared Stem Cell Facilities (CL1-00506-1,2), and CIRM Major Facilities (FA1-00617-1).

## Author contributions

S.A.S, G.A.L., and S.M.R. conceptualized the project. G.A.L. performed experiments and analyzed the results. S.A.S. and G.A.L. wrote the manuscript and prepared the figures. S.M.R. reviewed and gave feedback on the manuscript. Z.W, S.L, C.J.H, B.R.T, Z.W, S.K, S.A and E.W contributed to cloning, microscopy imaging ChIP-Seq and RNA-Seq experiments. N.B, E.M, and V.J contributed to ATAC-Seq and RNA-Seq analysis. J.W, N.H.T, T.UW and S.M.R performed and analyzed in vitro E2F1 nucleosome reconstitution and FP assays.

## Supplementary Data

**Supplementary Movie 1.** Live-cell imaging of EV-Clover, Fucci-CDT1-mCherry cells (+dox) for 24 hours with 20 minute image intervals. Left: Phase contrast. Middle: EV-Clover. Right: Fucci-CDT1-mCherry.

**Supplementary Movie 2.** Live-cell imaging of E2F1-Clover, Fucci-CDT1-mCherry cells (+dox) for 24 hours with 20 minute image intervals. Left: Phase contrast. Middle: E2F1-Clover. Right: Fucci-CDT1-mCherry.

**Supplementary Movie 3.** Live-cell imaging of EV-Clover, Fucci-CDT1-mCherry cells (+dox, +10% serum) for 24 hours with 20 minute image intervals. Left: Phase contrast. Middle: EV-Clover. Right: Fucci-CDT1-mCherry.

**Supplementary Movie 4.** Live-cell imaging of E2F12Clover, Fucci-CDT1-mCherry cells (+dox) for 24 hours with 20 minute image intervals. Left: Phase contrast. Middle: E2F2-Clover. Right: Fucci-CDT1-mCherry.

**Supplementary Movie 5.** Live-cell imaging of E2F1dDNA-Clover, Fucci-CDT1-mCherry cells (+dox) for 24 hours with 20 minute image intervals. Left: Phase contrast. Middle: E2F1dDNA-Clover. Right: Fucci-CDT1-mCherry.

**Supplementary Table 1**. Differential gene expression including overlap annotation between E2F1 and Serum treatment, showing overall basemean and log2 fold change per sample.

**Supplementary Table 2**. Full motif analysis results of serum-only and shared accessible promoter regions.

**Supplementary Table 3**. Integrated table with gene expression (RNA-Seq) showing overall basemean and Log2 fold change per sample, accessibility in promoter regions (ATAC-Seq) as measured by q-value for each peak region, and E2F1 binding in promoter regions (ChIP-Seq) as measured by q-value for each peak region.

**Supplementary Figure 1.**
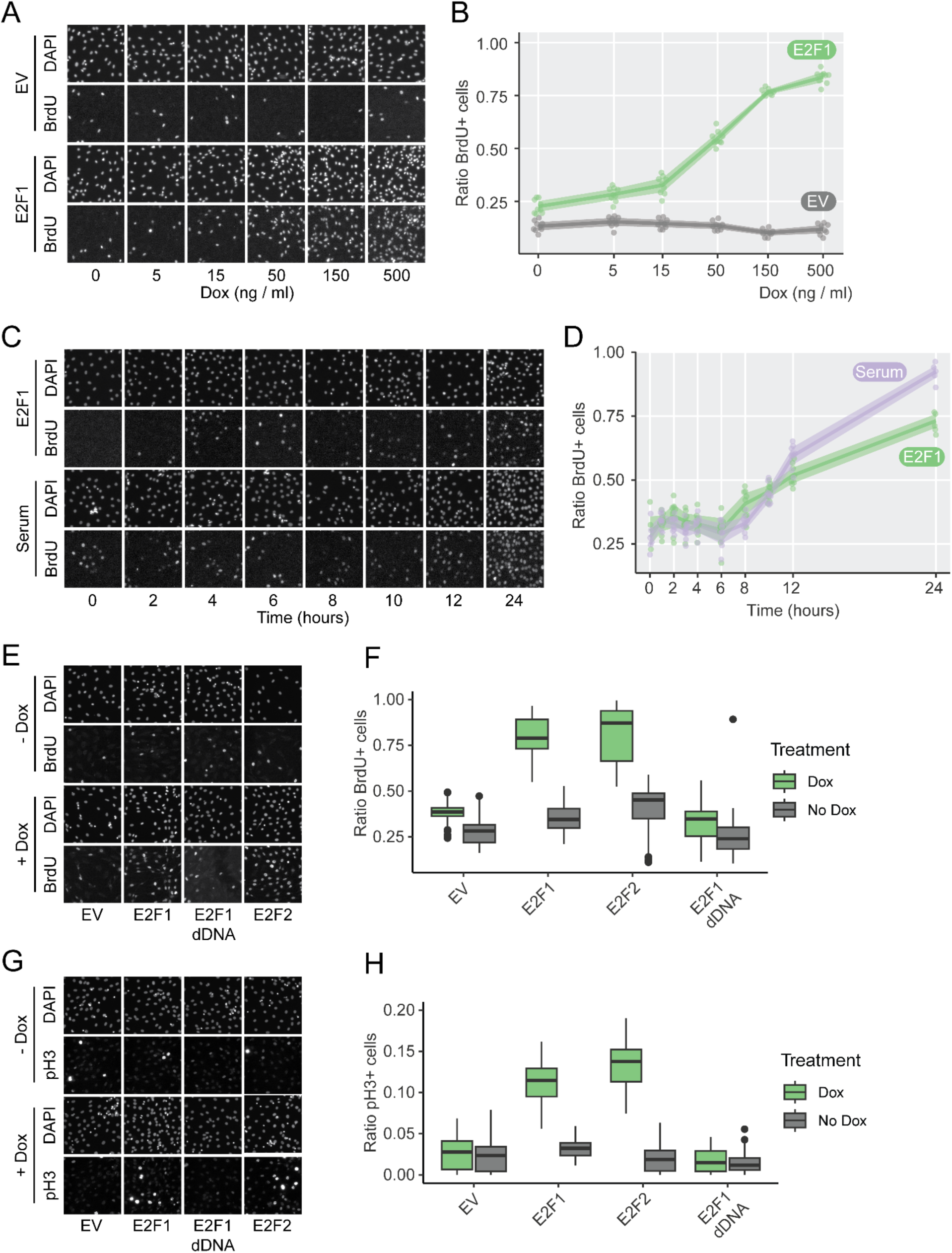
**related to Figure 1 and Figure 2. A)** Representative images of DAPI and BrdU staining after induction of E2F1 using increasing dox concentrations compared to EV control. **B)** Quantification of dose-dependent S-phase entry as measured by BrdU/DAPI ratio. **E)** Representative images of DAPI and BrdU staining to compare induction of EV, E2F1,, E2F2 and E2F1-dDNA under control (-dox) and induced (+dox) conditions. **F)** Quantification of S-phase entry as measured by BrdU/DAPI ratio, n = 3 independent experiments with 12-16 images each. **G)** Representative images of DAPI and pH3 staining to compare induction of EV, E2F1, E2F2 and E2F1-dDNA under control (-dox) and induced (+dox) conditions. **H)** Quantification of M-phase entry as measured by pH3/DAPI ratio, n = 3 independent experiments with 12-16 images each.

**Supplementary Figure 2.**
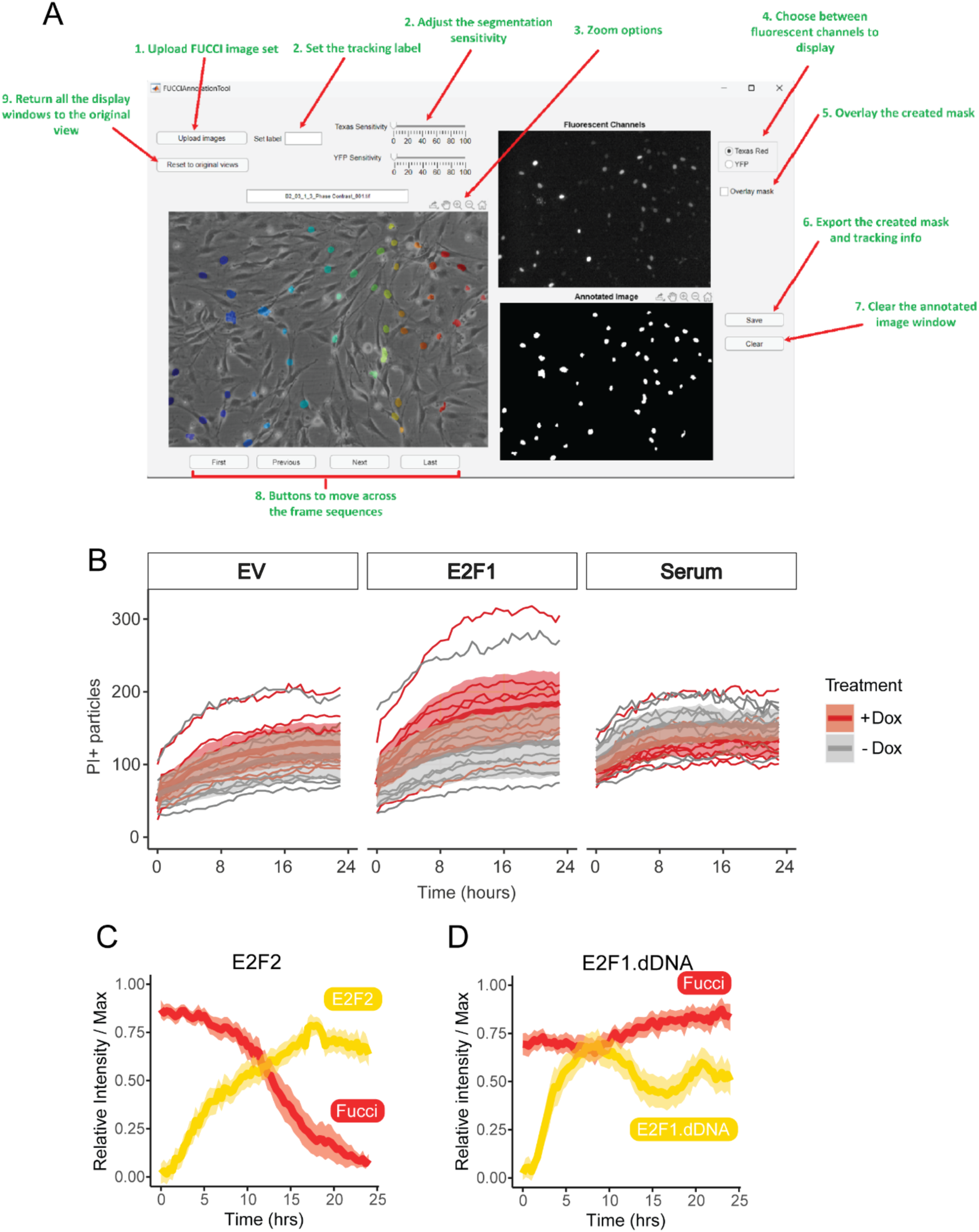
**related to Figure 2. A)** Interface of semi-automated live cell imaging quantification analysis software used. **B)** Single cell quantification of dox-treated E2F2-Clover::CDT1-mCherry cells (mean, 95% c.i., n = 73 individual cells). **C)** Single cell quantification of dox-treated E2F1-dDNA::CDT1-mCherry cells (mean, 95% c.i., n = 73 individual cells). **D)** Quantification of PI staining in live cell assays comparing EV control, E2F1 and serum treated cells, n = 9 image areas per condition.

**Supplementary Figure 3.**
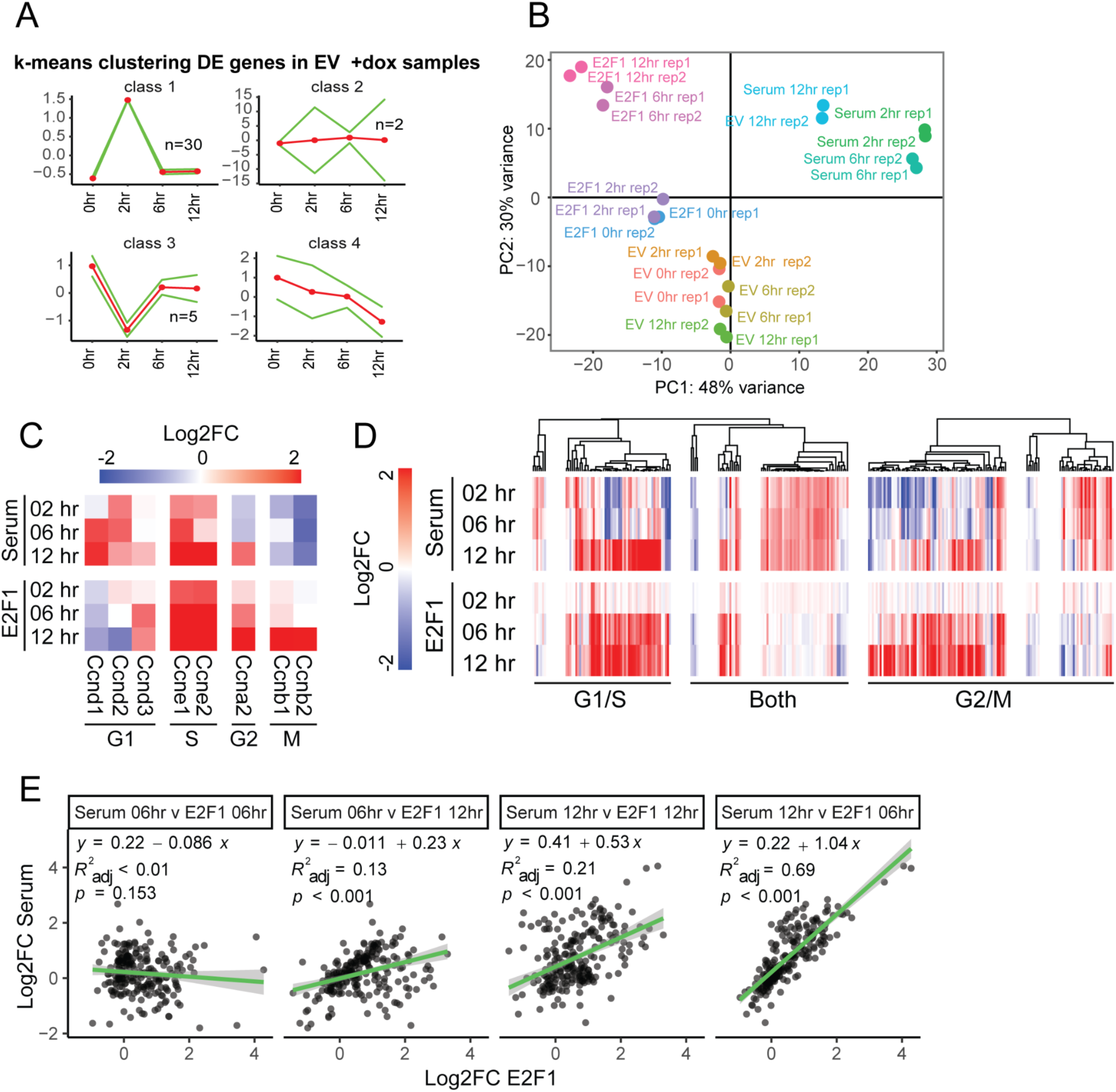
**related to Figure 3. A)** K-means clustering of EV differentially expressed genes. **B)** PCA plot of all RNA-seq samples. **C)** Heatmap indicating Log2FC of cell-cycle genes compared to 0 h**. D)** Heatmap indicating Log2FC of Reactome cell-cycle genes involved in G1/S or G2/M transition. Some genes were involved in G1/S and G2/M (Both). **E)** Linear regression showing pairwise comparisons of cell-cycle gene expression of 6h and 12h timepoints of serum treated and E2F1 treated cells.

**Supplementary Figure 4.**
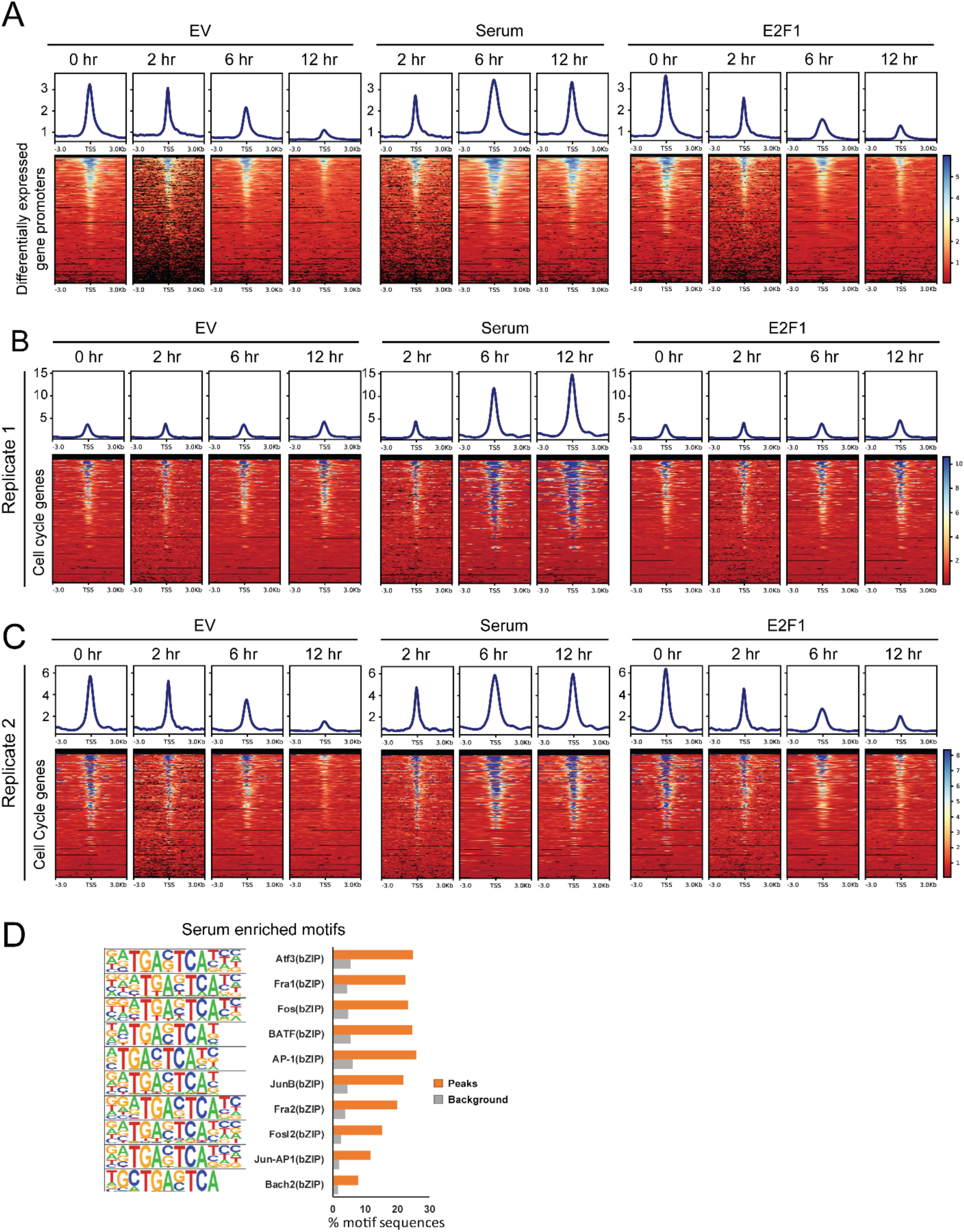
**related to Figure 4. A)** Replicate 2 of ATAC-Seq data showing accessibility profiles in promoters of differentially expressed genes **B)** ATAC-Seq data showing accessibility profiles in promoters of cell-cycle genes for Replicate 1, and **C)** for Replicate 2. **D)** Motif enrichment analysis of differentially accessible peaks enriched in serum mediated cell-cycle reentry showing top 10 motifs.

**Supplementary Figure 5.**
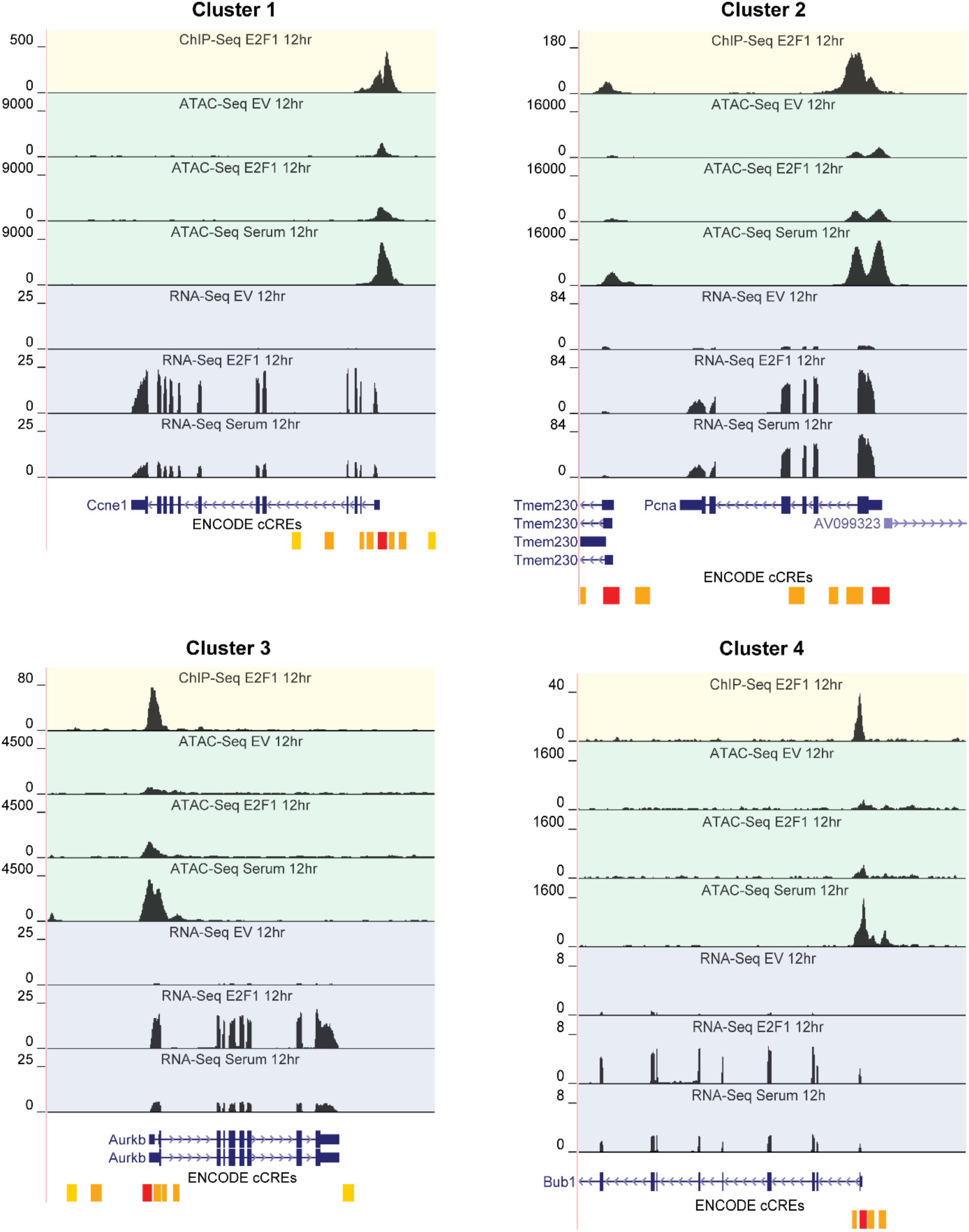
**related to Figure 5.** Multi-omic data visualized on the UCSC Genome Browser showing E2F1 binding (ChIP-Seq), chromatin accessibility (ATAC-Seq) and gene expression (RNA-Seq) for Ccne1 (Cluster 1), Pcna (Cluster 2), Aurkb (Cluster 3) and Bub1 (Cluster 4) loci. Cluster numbers indicate different chromatin accessibility levels as shown in Figure 5D.

## Notes

### Competing Interest Statement

The authors have declared no competing interest.

